# Segregational drift hinders the evolution of antibiotic resistance on polyploid replicons

**DOI:** 10.1101/2023.02.01.526651

**Authors:** Ana Garoña, Mario Santer, Nils F. Hülter, Hildegard Uecker, Tal Dagan

## Abstract

The emergence of antibiotic resistance under treatment depends on the availability of resistance alleles and their establishment in the population. Novel resistance alleles are encoded either in chromosomal or extrachromosomal genetic elements; both types may be present in multiple copies within the cell. However, the effect of polyploidy on the emergence of antibiotic resistance remains understudied. Here we show that the establishment of resistance alleles in microbial populations depends on the ploidy level. Evolving bacterial populations under selection for antibiotic resistance, we demonstrate that resistance alleles in polyploid elements are lost frequently in comparison to alleles in monoploid elements due to segregational drift. Integrating the experiments with a mathematical model, we find an agreement between the theoretical and empirical allele dynamics, confirming our understanding of the allele segregation process. Using the mathematical model, we further show that the effect of polyploidy on the establishment probability of beneficial alleles is strongest for low replicon copy numbers and plateaus for high replicon copy numbers. Our results suggest that the distribution of fitness effects for mutations that get fixed in a population depends on the replicon ploidy level. Our study indicates that strategies for drug treatment of bacterial infections should take into consideration the pathogen ploidy level.

## Introduction

Bacterial adaptation to novel environmental conditions depends on the availability of beneficial alleles and the dynamics of their proliferation within the population. However, novel alleles are initially rare in the population and therefore – even if they are adaptive – prone to stochastic loss due to genetic drift [1]. The probability that beneficial alleles survive the early proliferation phase and become established is termed ’establishment probability’. The probability of beneficial allele establishment (and eventually fixation) is a key concept in theoretical population genetics [2,3], dating back to the very early days of the field [1]. Understanding the determinants of allele establishment in bacterial populations is pivotal in the context of antibiotic resistance evolution. Antibiotic treatment confers a growth advantage to bacteria carrying resistance alleles, which might establish and subsequently rise to high numbers. The emergence of novel resistant strains of diverse human and livestock pathogens due to antibiotic treatment has worrisome consequences for global human health [4]. Sustainable treatment strategies are needed [5], for the development of which a profound understanding of the dynamics of beneficial alleles – including the early stochastic phase – is necessary. Several recent studies have supplied empirical evidence for stochastic loss of adaptive antibiotic resistance genotypes and factors that may be involved in their establishment [6–8]. However, the effect of bacterial ploidy on the establishment of antibiotic resistance remains understudied.

Prokaryotic genomes comprise chromosomes and extrachromosomal genetic elements, e.g., plasmids, collectively termed replicons. The number of chromosome copies in the cell is tightly controlled during replication and is synchronised with cell division [9,10]. Several prokaryotic taxa are known to harbour polyploid chromosomes. This includes cyanobacteria, e.g., *Synechococcus elongatus* that has a chromosome copy number of 3-4 [10,11]. Other examples are the human pathogens *Borrelia hermsii* and *B. burgdorferi*, which have a chromosome copy number of 4-14 and 10-20, respectively [12]. The number of chromosome copies in the cell may furthermore depend on the growth phase and nutrient conditions (e.g., refs. [13,14]); indeed, also seemingly monoploid bacteria, such as *E. coli*, can be polyploid during exponential growth [15,16]. The number of plasmid copies in a host cell depends on the plasmid type and the host genetics. Low copy number plasmids are found in 1-5 copies in the cell, while high copy number plasmids may reach 200 copies in the cell [17,18]. Similarly to chromosomes, the plasmid copy number may vary depending on the growth conditions [19,20].

The presence of multiple replicon copies in the cell has important implications for bacterial evolution. Experimental evolution of bacteria has shown that the mutational supply is positively associated with the replicon copy number, i.e., the probability of novel mutations can be higher for polyploid replicons (as demonstrated for multicopy plasmids in ref. [21]). Polyploidy furthermore allows for intracellular genetic diversity, also termed heterozygosity [22–24]. Alleles that emerge in monoploid chromosomes are inevitably inherited to daughter cells during cell division. In contrast, the segregation of alleles emerging in one out of many copies of a polyploid replicon is a neutral process and it may be unbalanced, such that only one daughter cell inherits the novel allele. The effect of genetic drift on novel plasmid alleles during cell division has been termed segregational drift [25]. The dynamics of alleles encoded in polyploid chromosomes (and multicopy plasmids) are thus affected by processes at two hierarchical levels: within the cell and within the population [25–29]. The mechanism of replication and the mode of segregation are important determinants of the allele dynamics on polyploid replicons [28,29]. A recent model on the fixation process of beneficial allele dynamics shows that heterozygous cells can persist for an extended period of time during the fixation process of high copy number replicons [29]. The coexistence of multiple plasmid alleles within a cell enables the host population to survive in fluctuating environmental conditions, e.g., under alternating drug pressures selecting for multiple antibiotic resistances [23]. The replicon copy number has furthermore important implications for the dynamics of rare novel alleles. Theory predicts that segregational drift will decrease the establishment probability of beneficial alleles when encoded in multicopy plasmids, at least in the absence of gene dosage effects [25,27]. Nonetheless, empirical evidence for the workings of segregational drift on the dynamics of beneficial alleles remains lacking.

Here we investigate the dynamics of a novel antibiotic resistance allele in an experimentally evolving bacterial population of *Acinetobacter baylyi* and compare the results between populations where the allele is encoded either on a polyploid (multicopy) plasmid or on a monoploid chromosome. We furthermore combine the experimental results with a mathematical model of bacterial population genetics to probe our mechanistic understanding of the dynamics of alleles on polyploid replicons.

## Results

### Rare alleles in multicopy replicons are prone to rapid loss

To experimentally quantify the effect of the replicon copy number on the fate of beneficial alleles, we studied the dynamics of an allele conferring resistance to the antibiotic kanamycin (*nptII*). The beneficial allele was introduced either on a polyploid (multicopy) plasmid or a monoploid chromosome in the model organism *A. baylyi*. Performing an experimental evolution experiment with serial transfers, we examined the effects of the initial allele frequency, the strength of the population bottlenecks, and the selection regime on allele dynamics (Fig 1a). We compare the results for three initial frequencies of cells containing the novel allele: *f*_0_ = 10^−4^ (ca. 10^5^ cells, high), *f*_0_ = 10^−6^ (ca. 10^3^ cells, moderate) and *f*_0_ = 10^−7^ (ca. 10^2^ cells, low). The control of the initial frequency of cells containing the novel plasmid allele was achieved via natural transformation employing varying donor DNA concentrations (700, 7, or 0.7 ng/μl DNA). The novel *nptII* allele is introduced into an ancestral population that carries the model plasmid pTAD-R, which has a copy number of ca. 15. Homologous recombination between the donor DNA and the ancestral plasmid creates heterozygous cells carrying both the ancestral and the novel alleles, where the intracellular frequency of the novel allele is one plasmid copy out of 15 due to the one-hit kinetics of natural transformation (Fig 1a; ref. [28]). For the chromosomal allele, we controlled the initial frequency by mixing strains carrying either the novel or the ancestral allele at the abovementioned frequencies. The evolved populations were serially transferred using two population bottleneck sizes: 1:100 (*b* = 10^−2^; ca. 10^7^ cells, weak bottleneck) and 1:1000 (*b* = 10^−3^; ca. 10^6^ cells, strong bottleneck) for over ca. 200 generations (30 serial transfers). The experiment was conducted with six replicates for each experimental condition. In the absence of antibiotics, the novel *nptII* allele has no observable fitness effect (Fig S1), such that changes in allele frequencies in the population over time are attributed to neutral processes rather than fitness differences between genotypes in the population.

**Fig 1.**
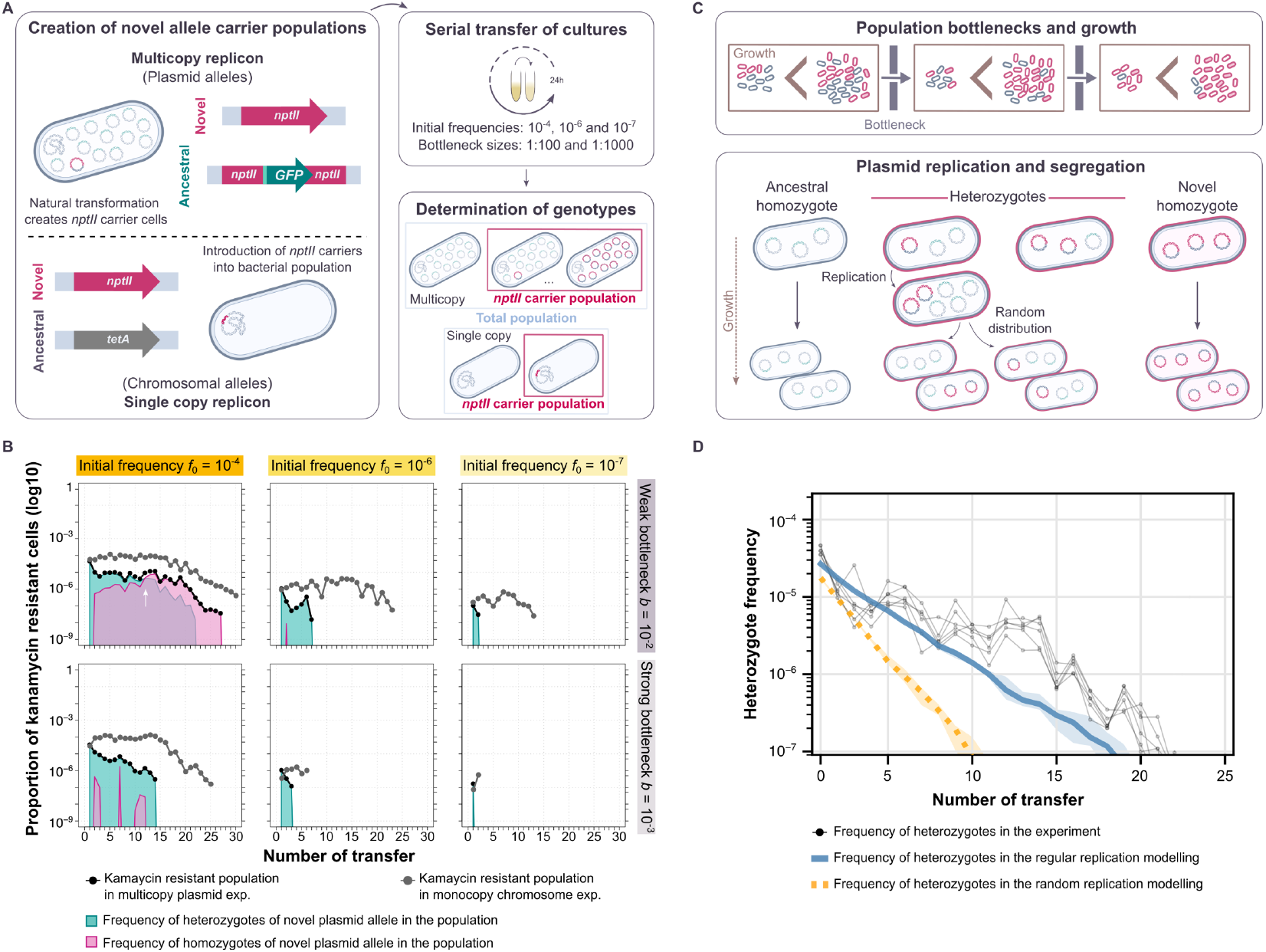
Integrating multicopy replicon allele dynamics *in vivo* and *in silico*. (A) Our experimental system enables us to create and follow a novel allele in bacterial populations over the course of an evolution experiment. The novel allele employed is a kanamycin resistance gene (*nptII*), which encodes aminoglycoside 3’-phosphotransferase II [42]. The ancestral plasmid allele is a green fluorescence protein (*gfp*) whose transcription is under the control of a LacI repressor [28]. The chromosomal ancestral allele is a tetracycline resistance gene (*tetA*). The bacterial populations were serially passaged under different conditions for approximately 200 generations. We followed the frequency of cells carrying the novel allele by selective plating on kanamycin-containing medium. For the plasmid replicon, we examined the genotypic composition of the populations by illuminating fluorescent colonies. (B) Allele dynamics of representative populations evolved under non-selective conditions. The white arrow marks the point at which the frequencies of heterozygous and novel homozygous cells are equal. The remaining experimental populations are shown in Fig S2 and S3. (C) Schematic diagram of the mathematical model of plasmid allele dynamics under periodic population bottlenecks (transfers). Bacterial populations grow until they reach a carrying capacity *N_c_* before each transfer. A proportion *b* of cells is randomly chosen for the next growth phase. During each growth phase, cells carrying only the ancestral allele (ancestral homozygotes) replicate at a rate that is *1 - s* times smaller than the rate of cells carrying the novel allele (heterozygotes and novel homozygotes). (D) Comparison between the allele dynamics in evolution experiments using the pTAD-R plasmid and in model simulations with replicon copy number *n*_rc_ = 15 under non-selective conditions (*s* = 0) for high initial frequencies, *f*_0_ = 10^−4^, and weak bottlenecks, *b* = 10^−2^. Bold lines show the median of the heterozygote frequencies before each transfer from 10 simulations for both modes of replication; shaded areas show the corresponding interquartile ranges. Thin lines show the heterozygote frequency from the six replicates of the evolution experiment. Simulation results were obtained by integrating the differential equations for bacterial growth and sampling new populations at each transfer from a multinomial distribution defined by the genotype frequencies at carrying capacity.

To demonstrate the effects of the initial frequency and the bottleneck strength, we first present the observed allele dynamics under non-selective conditions for a set of representative populations (Fig 1b). Our results show that with a high initial frequency (*f*_0_ = 10^−4^) and a weak bottleneck (*b* = 10^−2^), the dynamics of the novel allele in both plasmid and chromosome are characterized by a stable persistence during the first ca. 20 transfers, followed by a decrease in the fraction of kanamycin resistant cells. An eventual decrease in the frequency of the rare novel allele in the population is to be expected under genetic drift since the fixation probability corresponds to the initial allele frequency and is thus very low [30]. Notably, the frequency of kanamycin resistant cells decreases faster if the allele is on the multicopy plasmid than if it is on the chromosome. At the end of the experiment, the novel plasmid allele goes extinct, while the chromosomal allele is present in the population at a frequency of ca. 5×10^−7^.

Our system enables us to distinguish between heterozygous and homozygous cells for the novel plasmid allele. The subpopulation of cells that carries the novel allele initially consists of heterozygotes in which the novel allele is present in only one plasmid copy per cell. As the allele segregation takes place, kanamycin resistant homozygous cells arise and increase in frequency over time, until heterozygous cells are no longer present and intracellular genetic diversity is lost. Our results further show that when the initial frequency of the cells carrying the novel allele is lower, the novel allele is maintained in the population for shorter times (Fig 1b; *f*_0_ = 10^−6^ and *f*_0_ = 10^−7^). The bottleneck size applied during the experiment has a significant effect on the allele dynamics, which are characterized by a rapid extinction in the populations transferred with a strong bottleneck (*b* = 10^−3^) compared to those populations evolved with a weak bottleneck. To sum up, under non-selective conditions, both plasmid and chromosomal alleles decrease over time and eventually are lost in most populations by the end of the experiment. Nonetheless, the loss of the plasmid allele is faster in all tested combinations of initial frequencies and population bottlenecks. Our results are in agreement with previous observations on the effect of segregational drift on the dynamics of alleles encoded in polyploid replicons [25,27,28].

### Mathematical modelling predicts the plasmid replication mode from the empirical plasmid allele dynamics

To formulate the theoretical expectation for the allele dynamics in the experiments, we adapted a previous mathematical model of vertical replicon inheritance in bacterial populations to match the experimental design of the evolution experiment (Fig 1c; refs. [27,29]). The genotype of a cell is defined by the number of novel and ancestral replicon copies at cell birth, with the total number of novel replicon copies ranging between 0 and the replicon copy number *n*_rc_. Cells in the simulations of monoploid replicons (*n*_rc_ = 1), such as the chromosome in *A. baylyi*, harbor either the ancestral or the novel allele. Cells in the simulations of multicopy replicons (*n*_rc_ > 1), e.g., model plasmid pTAD-R, can be homozygous for the ancestral or for the novel allele, or heterozygous, carrying both allele variants in various proportions. Our model includes two possibilities for the mode of replication: under the *regular replication* mode, all *n*_rc_ replicon copies are duplicated before a cell divides. Under the *random replication* mode, single copies are selected randomly for replication in a repetitive process until the cell carries 2 x *n*_rc_ copies (refs. [27,29]; see Methods for details). For either mode of replication, the replicon copies are distributed randomly into the two daughter cells at cell division (similarly to pTAD-R), where each cell inherits *n*_rc_ copies.

The replication mode of pTAD-R backbone has not been examined experimentally so far. To determine which modelled mode of replication better matches the experimental system, we simulated the segregation dynamics of a novel allele carried by a polyploid replicon and compared the simulated dynamics to the experimental results. The simulation was performed for both replication modes under neutrality (*s* = 0) with a replicon having *n*_rc_ = 15, as model plasmid pTAD-R. In this simulation, we opted for the highest initial frequency, *f*_0_ = 10^−4^, and the weak bottleneck, *b* = 10^−2^, where stochastic effects other than segregational drift are weakest. The model results show that the frequency of heterozygous cells rapidly decreases at a constant rate. The comparison between the frequency of heterozygotes from simulations under the mode of regular replication and the experimental results shows a good qualitative agreement for the first 11 transfers (Fig 1d). From transfer 12 onwards, the model underestimates the frequency of heterozygous cells by up to ten-fold. Simulations under the mode of random replication, however, overestimate the decay of heterozygotes frequency already during the very first transfers. Consequently, we opted to use the regular replication mode for further simulations of pTAD-R allele dynamics. The correspondence between the simulation results under the regular replication mode and the allele dynamics in the evolution experiment suggests that the replication of plasmid pTAD-R is not random and may follow a regular mode of replication.

### Beneficial alleles on polyploid replicons are prone to loss due to segregational drift

To examine the effect of replicon polyploidy on the dynamics of beneficial alleles, we experimentally evolved the bacterial populations under conditions where the novel allele is beneficial, i.e., under selection for kanamycin resistance. In order to identify a comparable selection regime for both replicon types, we quantified the fitness effect of the *nptII* allele when encoded in the multicopy plasmid or single copy chromosome, employing increasing kanamycin concentrations (Fig S4 and S5). During the evolution experiment, we applied a constant antibiotic pressure, which confers cells that are homozygous for the novel allele a fitness benefit of ca. 10% (i.e., selection coefficient *s* = 0.1). Since the concentration of antibiotic employed to achieve a fitness benefit of ca. 10% in multicopy and single copy replicon types is highly similar, we conclude that the effect of gene dosage in our system is negligible. Note that the effect of selection is restricted to processes at the population level; antibiotic selection has no known effect on plasmid replication and segregation. We examined the effect of selection on the dynamics of the novel allele in combination with the effect of initial frequency and population bottleneck size. For each replicon type, we thus tested a total of 12 parameter combinations, each with 6 replicates (Fig 2 and Fig S2 and S3).

**Fig 2.**
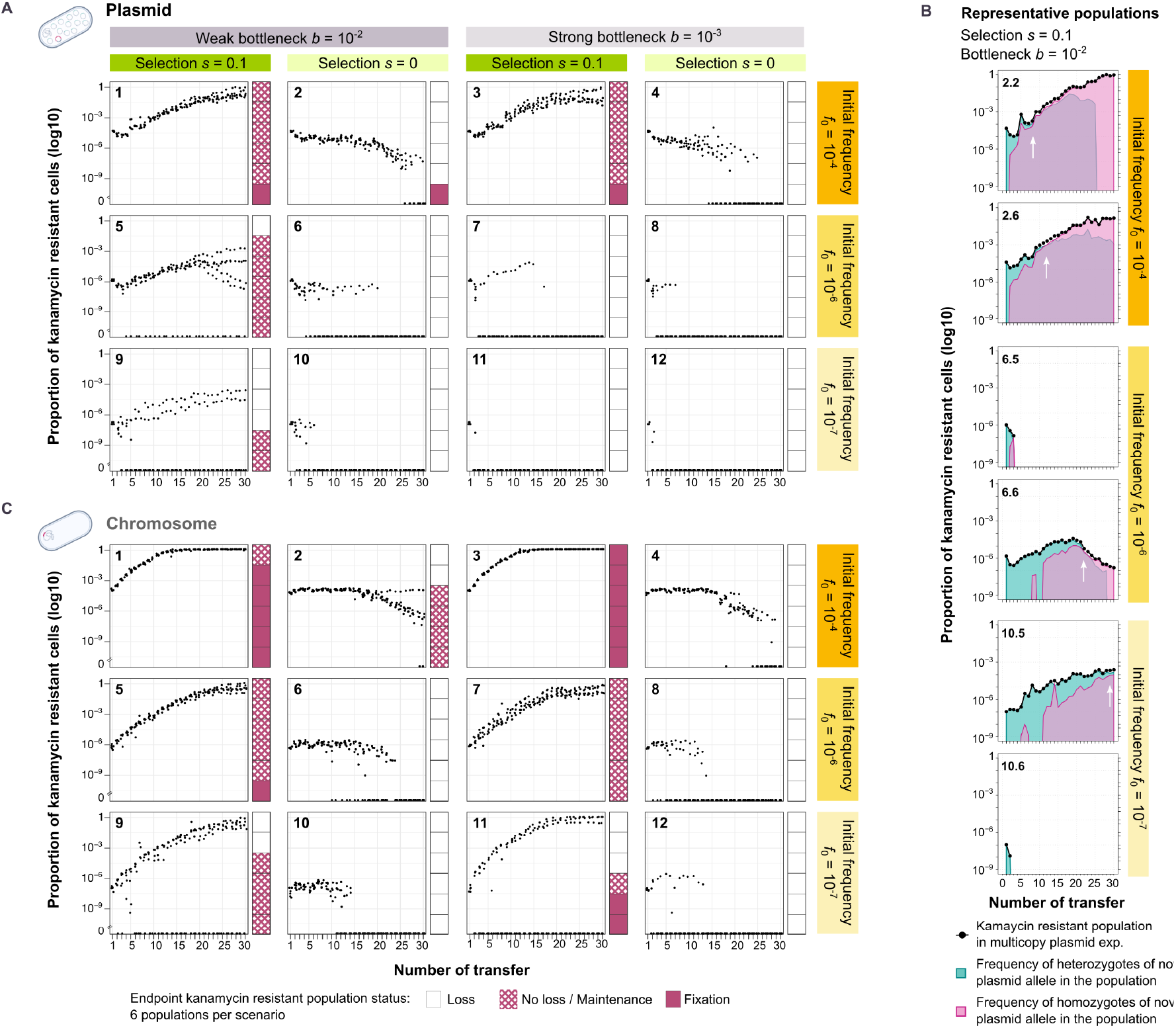
Plasmid and chromosomal allele dynamics under selective and non-selective conditions. (A) The dynamics of a plasmid allele over approximately 200 generations. Points correspond to the proportion of kanamycin resistant cells in each population and sampling time. Stacked bar plots present the state of the kanamycin resistant population at the end of the experiment. The fixation threshold is set at a frequency of 0.9. (B) Plasmid allele dynamics of representative populations under selective conditions and weak bottleneck. The white arrow marks the point at which the frequencies of heterozygous and novel homozygous cells are equal. The remaining experimental populations are shown in Fig S2 and S3). (C) The dynamics of a chromosomal allele over approximately 200 generations.

We first focus on the results for the plasmid encoded allele. As expected, the novel allele is maintained for longer and lost less often under selection than under neutral conditions. With high initial frequencies (*f*_0_ = 10^−4^), the fraction of kanamycin-resistant cells increased in the population over time, regardless of the population bottleneck (Fig 2a, plots 1,3). The frequency of cells that carry the novel allele at the end of the experiment is close to the fixation threshold of 90% in most replicates. In contrast, at low or moderate initial frequencies, the novel plasmid allele was quickly lost in most populations (Fig 2a). In populations that evolved under weak bottlenecks, the novel plasmid allele is lost in 1 out of 6 replicates for moderate initial frequencies and in 4 out of 6 replicates for low initial frequencies (Fig 2a, plots 5,9). In some replicate populations where the allele is maintained, we observed a slow decrease in the fraction of kanamycin-resistant cells from the 20^th^ transfer on. Under the strong bottleneck, none of the replicate populations maintained the novel allele for moderate or low initial frequencies (Fig 2a, plots 7,11).

The combination of selection and initial frequency has an effect also on the genotypic composition of the resistant cell population. For high initial frequency (*f*_0_ = 10^−4^) under selective conditions, the frequency of cells that are homozygous for the novel allele increases rapidly and reaches a similar frequency to that of the heterozygotes within ca. 5-10 transfers. This initial rise in frequency is faster in comparison to that observed under non-selective conditions (ca. 10-15 transfers; Fig 1b). Under selection both genotypes are maintained in the population for a longer period of time; the heterozygotes decrease in frequency and are eventually lost from the population in a slower process (from ca. 25th onwards) (Fig 2b; Fig S2 and S3) in comparison to non-selective conditions. With medium and low initial frequencies, the homozygotes frequency increases slower, such that similar frequencies of homozygotes and heterozygotes are observed later in the experiment (Fig 2b). Under selective conditions, both resistant genotypes are maintained for a longer time, where the relative frequencies of homozygotes and heterozygotes are the manifestation of segregational drift.

To further examine the effect of polyploidy on novel allele dynamics we compared the dynamics observed for the plasmid-encoded allele to those observed for the chromosome-encoded allele. When the allele is encoded in the plasmid, the kanamycin resistant cell fraction increases slower and the time to fixation is longer in comparison to the chromosome-encoded allele (Fig 2c). The slower increase of the kanamycin resistant cell frequency in the experiments with the plasmid encoded allele is best observed at mid-experiment (ca. transfer 15) for the high initial frequency (*f*_0_ = 10^−4^), where the frequency of kanamycin resistant cells is 100 times lower in comparison to the chromosome experiment. At the end of the experiment, the kanamycin resistance phenotype was fixed in a single population within the plasmid experiment, in contrast to chromosome experiment where the resistance phenotype was fixed in the majority of populations (Fig 2a, c, plots 1,3). At lower initial frequencies (*f*_0_ = 10^−6^, *f*_0_ = 10^−7^), the differences between the frequency of kanamycin resistant cells observed in the plasmid and chromosome experiments are even more pronounced, with up to ca. 1000-fold in favour of the chromosome populations (Fig 2a, c, plot 5). The most striking difference between plasmid and chromosome experiments is observed in populations evolved under a strong bottleneck (*b* = 10^−3^). In those populations the plasmid-encoded allele is lost in all replicate populations (Fig 2a, c, plots 7,11), in contrast to the chromosome experiment where the allele is maintained. To conclude, the opposing effect of segregational drift and selection lead to longer fixation time and loss of plasmid alleles in comparison to chromosomal alleles. Our results demonstrate that segregational drift has a pivotal role in the evolution of polyploid replicons.

An alternative explanation for the observed loss of the novel beneficial allele could be the adaptation of the originally kanamycin-sensitive cells to the antibiotic environment, such that the novel allele is no longer beneficial. To rule out this alternative, we performed a competition experiment where the ancestral kanamycin-resistant populations were competed against the evolved kanamycin sensitive populations (Fig S6). Our results show that the evolved kanamycin sensitive populations still had a lower fitness under selective conditions compared to the ancestral kanamycin-resistant populations. Consequently, we conclude that the dynamics of the novel plasmid allele observed during the evolution experiment are the result of plasmid allele segregation rather than adaptation of the kanamycin sensitive population to kanamycin.

### The negative effect of segregational drift on the fixation of the beneficial allele is in agreement with model simulations

To examine whether the previous theoretical predictions on the effect of replicon polyploidy are validated by the empirical results from the experimental evolution experiment, we performed further simulations under selective conditions. In our model, we assume that cells carrying only the ancestral allele replicate at a rate 1 – *s*, and cells carrying at least one copy of the novel allele replicate at a rate 1. The model assumes no effect of gene dosage.

Since the implementation of the model parameter *s* varies from the experimental determination of selection regime, we first estimated the parameter *s* from the mean frequencies of the kanamycin resistant sub-population. The *s* values yielding an optimal fit between the simulated dynamics of the novel allele and the corresponding dynamics in the evolution experiment for large initial frequencies, *f*_0_ = 10^−4^, and weak bottlenecks, *b* = 10^−2^, were selected for further simulations (Fig S7). Those deviated slightly from the experimental assessment of the fitness benefit of the novel allele: *s* = 0.086 for the polyploid replicon (*n*_rc_ = 15) and *s* = 0.14 for the monoploid replicon (*n*_rc_ = 1).

To examine how well our model captures the experimental results, we first performed simulations that correspond to the experimental conditions (Fig 3). The dynamics of the novel phenotype (kanamycin resistance) were overall in agreement with the results of the evolution experiment for both replicon types, albeit the frequency of the novel phenotype is slightly overestimated by the model given low or moderate initial frequencies (*f*_0_ = 10^−4^ or 10^−6^) or strong bottlenecks, *b* = 10^−3^. The simulation results for the polyploid replicon show that the rise in frequency of heterozygotes in the experiment is well captured by the model simulations, for high and moderate initial frequencies (*f*_0_ = 10^− 4^ and 10^−6^) and bottleneck size combinations. Nonetheless, the frequency of heterozygotes as observed in the evolution experiment is underestimated by the model. The frequency of heterozygotes in the simulation starts decaying at nearly the same transfer as in the experiment, yet heterozygotes in the evolution experiment reach frequencies that are 10- to 100-fold higher in comparison to the simulations. The overestimated heterozygous loss rate in the simulation could possibly be explained by a (temporarily) increased plasmid copy number during the evolution experiment. Note that a comparison of the simulated allele dynamics to the plasmid experiment for the low initial frequency (*f*_0_ = 10^−7^) is not possible as most replicates in those experiments went extinct. Using the model simulation for the low initial frequency thus supplies a theoretical expectation for the beneficial allele dynamics in rare cases where the novel allele may persist in the population. In our theoretical model, we describe population growth between bottlenecks deterministically, and bottlenecks are thus the only possible source of allele loss.

**Fig 3.**
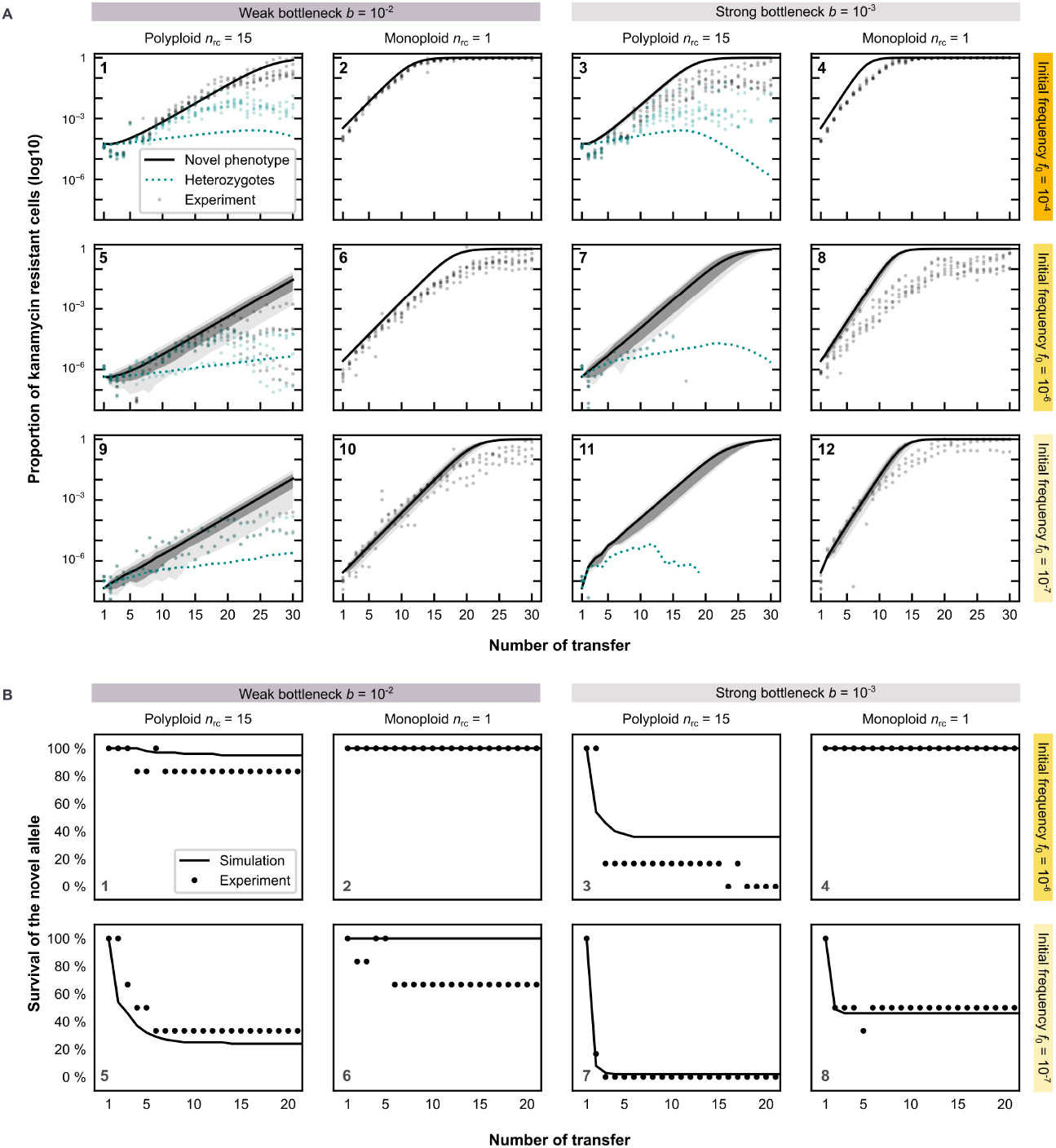
Model simulations of beneficial allele dynamics for polyploid (*n*_rc_ = 15) and monoploid replicons. (A) Cell frequencies of the novel phenotype (parallel to kanamycin resistant cells) (solid) and heterozygous cells (dotted) at the end of each population transfer obtained from 100 simulations for a replicon with copy number *n*_rc_ = 15 (1,3,5; for the multicopy plasmid) and with *n*_rc_ = 1 (2,4,6; reflecting the chromosome). Lines show the mean of positive frequencies over the number of transfers, i.e., the average proportion of cells having the novel phenotype from all simulation trajectories, where the novel allele has not (yet) gone extinct. The initial population at the start of the first transfer consists of ancestral cells and a small proportion, *f*, of cells with one novel replicon copy, where *f*_0_ = 10^−4^ in (1-4), *f*_0_ = 10^−6^ in (5-8), and *f*_0_ = 10^−7^ in (8-12). The grey (dark grey) areas show the 99% (67%) confidence intervals of positive mutant cell frequencies. The green dotted lines show the mean frequency of heterozygous cells carrying both the ancestral and novel plasmid allele (conditioned on positive frequencies). Grey and green markers show the frequencies of kanamycin resistant cells and heterozygous cells, respectively, of the corresponding experimental replicates. (B) Survival of the novel allele over number of transfers in model simulations (lines) and in the evolution experiment (markers). The fraction of kanamycin resistant cells of 100 simulation trajectories and 6 experimental replicates, respectively, is shown for each parameter combination. Simulation results were obtained by integrating Eq. (A11; in ref. [29]) until the population size has reached *N*_*c*_(*t*) cells in each growth phase (transfer), where *N*_*c*_(*t*) is taken from the mean experimental carrying capacity of each transfer *t*. A population bottleneck is applied after each growth phase, where a proportion of *b* = 10^−2^ or *b* = 10^−3^ cells are randomly selected from a multinomial distribution defined by the genotype frequencies at the end of the growth phase.

The simulation results for moderate and low initial frequencies (*f*_0_ = 10^−6^ and 10^−7^) predict a higher variability in the kanamycin resistant cell frequencies per transfer for the polyploid replicon in comparison to the monoploid replicon (Fig 3a, plots 5-12). This prediction conforms to the results of the evolution experiments (Fig S8).

We further compare the survival probability of the novel allele over time with the experimental results for the fraction of populations in which the allele is still present after a given transfer (Fig 3b). The model shows a remarkable agreement with the experimental results, capturing the main trends and often even making quantitatively accurate predictions.

### The establishment probability of alleles on polyploid replicons depends on the strength of selection and the replicon copy number

So far, we simulated polyploid replicons with a copy number and a selection coefficient that reflect the pTAD plasmid copy number and the beneficial effect of the *nptII* allele. To further examine the influence of the replicon ploidy level on the survival of novel beneficial alleles, we performed simulations for a range of replicon copy numbers and selection coefficients. The simulation results show that the establishment probability of a beneficial allele, *p_est_*, decreases with the replicon copy number (Fig 4), as previously found in a fully stochastic model without bottlenecks [27]. The strength of selection, *s*^(50%)^, required for equal chances of allele loss and allele establishment, *p*_est_ = 50%, rises substantially within the range of copy number <5 and reaches a plateau for intermediate to high copy numbers. For stronger bottlenecks, *b* = 10^−3^, the threshold for the strength of selection, *s*^(50%)^, is shifted to higher values of s, while the functional form with respect to the influence of the replicon copy number, *n*, on *s*^(50%)^ is qualitatively conserved (Fig 4). For the weak bottleneck, the model predicts an *s*^(50%)^ that is equivalent to a beneficial allele having a positive fitness effect of ca 20%, while in the model of strong population bottlenecks, *s*^(50%)^ is equivalent to an allele having a positive fitness effect of ca 45% in comparison to the ancestral allele. Our model simulations thus demonstrate that the effect of replicon ploidy on the establishment probability of beneficial alleles is particularly pronounced at low replicon numbers. The range of <10 copy number is frequently observed in broad host range plasmids and is also typically reached by chromosomes of polyploid bacteria and of seemingly monoploid organisms during their growth phase (e.g., *E. coli* refs. [16,26,31]).

**Fig 4.**
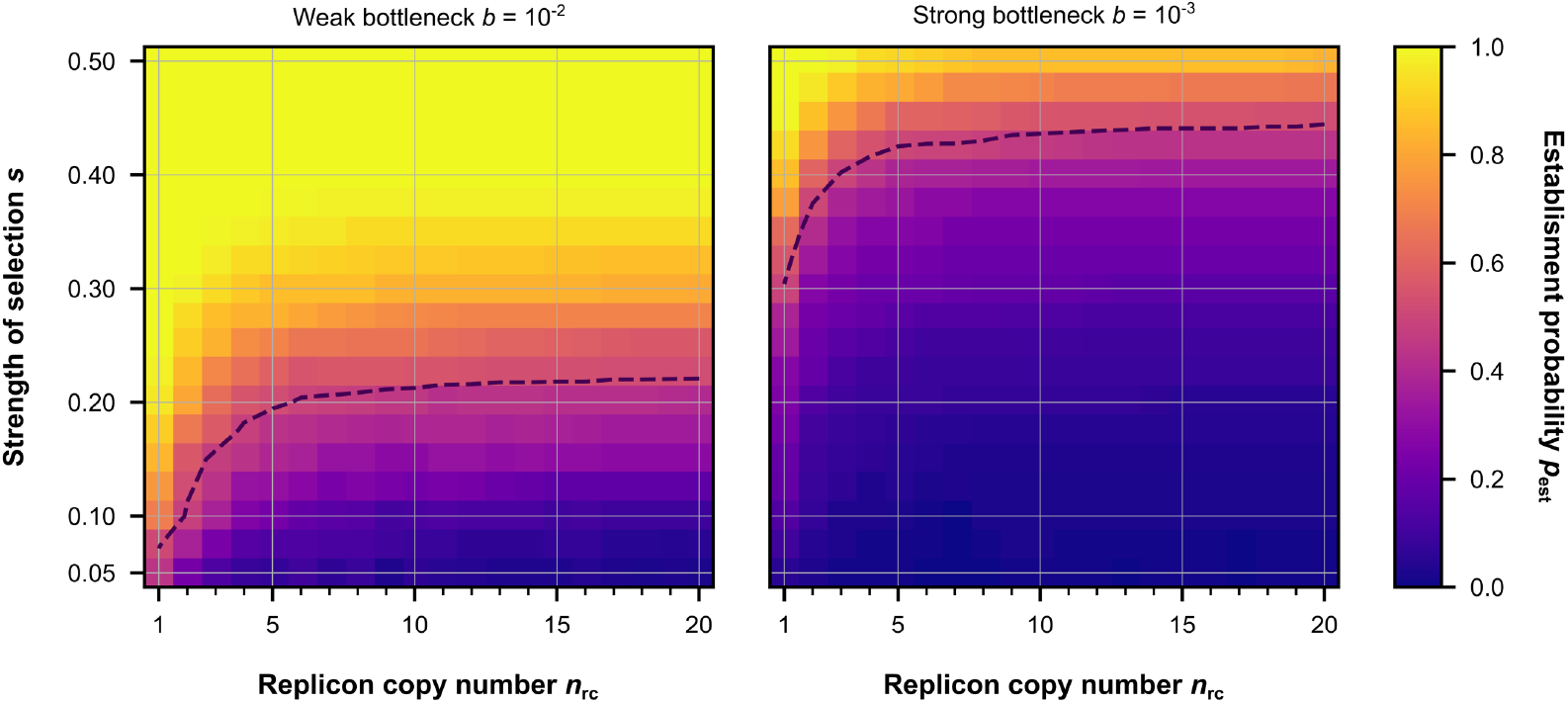
The influence of the replicon copy number (*n*_rc_) and the strength of selection (*s*) on the establishment probability of a beneficial allele. Allele establishment is achieved if the sub-population of cells with the novel phenotype is sufficiently large, such that stochastic loss of the allele becomes unlikely. Allele establishment in the simulation is defined by the time point when the frequency of novel homozygotes at the end of a growth phase is sufficiently high such that the probability of allele extinction is below 10^−4^. The initial population consists of 10^7^ cells, where one cell has a single mutant replicon copy and the rest is of the wild-type. The cell population grows between until a carrying capacity *N*_c_ = 10^9^ before a population bottleneck occurs, where a fraction *b* = 10^−2^ or *b* = 10^−3^ of cells is transferred to form the new population in the next growth phase. Simulations stop either when the novel allele goes extinct or when the allele is established in the population.

## Discussion

Polyploidy allows the coexistence of multiple alleles on a replicon – chromosome or plasmid – within a single cell, thereby creating intracellular genetic diversity. The establishment of alleles encoded in polyploid replicons is thus affected by drift and selection at two hierarchical levels: the collective of replicons within a cell and the collective of cells within the population. Intracellular processes – replication and segregation – generate an effect of genetic drift that interferes with the establishment of novel alleles.

A key aspect influencing the dynamics of novel alleles in polyploid replicons is the mode of their replication and its coordination with cell division [9,29,32]. Regular replication is a deterministic mode of replication, where each replicon copy is duplicated exactly once prior to cell division. In contrast, in random replication, replicon copies are randomly selected for replication throughout the cell growth cycle [33]. Using the regular replication mode, our model yielded accurate predictions of allele segregation rates and the decrease in heterozygote frequency in the early phase of the evolution experiment under non-selective conditions. Employing our mathematical model, we deduce that our model plasmid pTAD-R may have a regular mode of replication. Notably, an experimental determination of the replication mode is a laborious endeavour, with only few studies that supplied a characterisation of specific replicons, including plasmids [34–36] and chromosomes [37]. Our study thus presents a potential novel approach for the inference of replication mode using a combination of experimentally determined and mathematically modelled allele dynamics, which could be applied to other replicons.

Rare alleles are subject to a high risk of stochastic loss, which was predicted early on by Haldane’s theoretical result on the establishment probability, *2*s**, for mutations encoded in monoploid replicon under selection of strength **s** (assuming a Poisson offspring number) [1]. For polyploid replicons, the presence of intracellular genetic diversity creates a sub-population structure that can affect the establishment process of novel alleles. In our setup, the novel allele is initially introduced in a single copy out of 15 replicons within the cell. Thus, the relative frequency of novel alleles in polyploid replicons is smaller in comparison to alleles in monoploid replicons (*n*_rc_ > 1). Loss of the novel allele is highly unlikely once the allele reaches a sufficiently high frequency in the population, as in the results of our high initial frequency scenario (10^−4^) under selection. However, novel alleles emerging via *de novo* mutations or recombination are expected to have a lower initial frequency (similar to our experimental frequencies 10^−6^ and 10^−7^). The establishment of such low frequency alleles would require an escape from stochastic loss in order to rise to high frequencies, which is hindered in polyploid replicons by the decreased total relative allele frequency and segregational drift. At the same time, the mutational supply is higher in polyploid replicons [21], which may outweigh the effect of segregational drift, depending on the degree of dominance of beneficial alleles [27]. Our current results show that rare beneficial alleles are more prone to stochastic loss and rise slower to fixation if the allele is present on a multicopy replicon (i.e., the pTAD-R plasmid) compared to alleles on a single-copy replicon (i.e., *A. baylyi* chromosome). A comparison of the experimental and mathematical results confirms that theory on establishment probabilities, as approximated by survival probabilities (starting from low initial frequencies), is highly suitable to capture and predict the empirical observations.

Comparing the effects of the selection regime and the replicon copy number in our theoretical model, we found that the minimal strength of selection to reach an establishment probability of 50% (*p*_est_ = 50%) increases with the replicon copy number. The establishment probability *p*_est_ is most sensitive to the replicon copy number in a range of 1-10 copies per cell, which coincides with the copy number range of common polyploid chromosomes and natural plasmids. The effect of the replicon copy number has special significance in the context of antibiotic resistance evolution. Our results suggest that novel antibiotic resistance variants will follow different evolutionary trajectories depending on the type of replicon where they emerge, i.e., chromosomes, low copy number plasmids or medium to high copy number plasmids. We hypothesise that the dependency of the establishment probability on the replicon copy number has an impact on the distribution of fitness effects of resistance mutations that reach fixation. In other words, applying drug doses that lead to rapid fixation of resistance alleles in monoploid replicons, may lead to loss of the resistance allele when the allele is encoded in a polyploid replicon. Taken together, segregational drift has an effect on the evolutionary rate of polyploid replicons by influencing the dynamics of beneficial alleles, from their emergence to their eventual loss or fixation.

## Materials and Methods

### Bacterial strains, plasmids and culture conditions

The *A. baylyi* strain BD413 (DSM No. 588, German Collection of Microorganisms and Cell Cultures, DSMZ) also known as strain ADP1 (GenBank accession no. NC_005966.1) was used as the model organism in all experiments. For the chromosomal allele experiments, *A. baylyi* strain BD4 was also used. Either *A. baylyi* BD413 or BD4 were used during plasmid or chromosomal modification constructions. Primers used in this study are listed in supplementary tables (Supplementary Table 1). *A. baylyi* was propagated at 30 °C in LB medium in liquid shaking cultures or plates. For molecular cloning and plating, growth limiting factors, such as antibiotics, for the selection of plasmid/mutation carrying cells were used at the following concentrations: kanamycin 10 μg per ml, tetracycline 5 μg per ml, sucrose 50g per l. IPTG (Isopropyl β- d-1-thiogalactopyranoside) was added to the media to a final concentration of 1 mM when derepression of the LacI-repressed p*Trc* promoter was desired. Plasmids were extracted using the GeneJET Plasmid Miniprep Kit (Thermo Fisher Scientific). DNA quantification was done using the Multiskan GO spectrophotometer instrument (Thermo Fisher Scientific). All construction steps were validated by either restriction enzyme digestion or Sanger sequencing. The proof-reading Phusion polymerase (Thermo Scientific) was used for all PCR-based cloning.

The model plasmid pTAD employed in this study was constructed previously [28). pTAD-R was introduced into a naïve *A. baylyi* strain BD413 (ADP1), where it is stably maintained due to a toxin-antitoxin system. The *A. baylyi* strains containing the chromosomal alleles were constructed via natural transformation followed by homologous recombination between an allele containing PCR product and the chromosome. The antibiotic resistance alleles were introduced independently in two *A. baylyi* strains in the same genomic region, the *alkM* gene (ACIAD_RS06515). The kanamycin resistance allele (*nptII*) was inserted in the chromosome of *A. baylyi* BD413 strain (auxotrophic for tryptophan). The kanamycin resistance ancestral allele (*tetA*) encoding a tetracycline efflux protein, was inserted in the chromosome of *A. baylyi* BD4 (tryptophan prototroph wild-type). The construction of the PCR product was done by SOEing PCR. Two homologous regions to the *A. baylyi* chromosome flank the antibiotic resistance gene and were amplified by primer pairs (alkM-1up-fw/alkM_2up-rv+o1) and (alkM_5down-fw+o2/alkM_6down-rv). The *sacB-nptII* was amplified from the gene targeting vector system pGT41 [38) using primer pair (sacB-fw+o1/nptII-rv+o2). The *tetA* gene was amplified from plasmid pRKNH3-ΔI a RK2 derivative [39) employing primer pair (ptetA_tetA_fw+o1/ tetA_rv+o2). The three fragments were joined in a SOEing PCR reaction.

### Fitness experiments

The relative fitness (*w*) of the ancestral allele carrying strain versus the novel allele carrying strain was estimated by direct pairwise competition experiments, with 8 replicate populations for the plasmid replicon and 6 replicate populations for the chromosomal replicon. The determination of the selective coefficient given by the selection treatment was calculated via determination of the relative fitness of the ancestral allele carrying strain versus the novel allele carrying strain in direct pairwise competition experiments under selective conditions with 6 replicates populations for both plasmid and chromosomal replicons. The relative fitness (*w*) of the host population at the beginning of the experiment vs the non-host population at the end of the experiment was estimated by direct pairwise competition experiments with 8 replicate populations per experimental condition tested (conditions 8 and 12). All competition experiments were initiated with a 1:1 ratio of the competing strains, diluted 1:100 from overnight cultures, pre-conditioned to growth in liquid media for two days. The relative fitness is calculated by calculating the growth of each of the competitors after 24 hours of growth, the cultures are serially transferred for 5 days and the populations determined. Serial transfer was done in non-selective media for ancestral carrying vs novel allele carrying population competition and on selective media (kanamycin 0.375 μg/ml) for selective coefficient determination and host vs non-host competition. The two competitors are distinguished by plating on non-selective (LB supplemented with IPTG 1mM) and selective media (LB supplemented with kanamycin 10 μg/ml for novel allele carrying strains (chromosome or plasmid) or M9 for ancestral allele carrying chromosomal strains).

### Natural transformation of *Acinetobacter baylyi*

The preparation of competent cells of *A. baylyi* carrying pTAD-R was prepared as described previously in ref. [38]. Briefly, the cells were grown at 30 °C overnight with shaking in 2 ml of LB medium and used for the inoculation (1:100) of fresh cultures. These cultures were grown until early stationary phase (approx. 1×10^9^ cells per ml), cooled down and then stored as concentrated stocks (1×10^10^ cell per ml) at −80 °C.

For each transformation performed in the study, the frozen competent cells were thawed on ice and diluted to a cell density of 2.5×10^8^ cells per ml. The donor DNA was a PCR fragment (for a detailed description see ref. [28]) containing the non-disrupted *nptII*^+^ as the single segment of sequence identity to the recipient plasmid. After 90 minutes of incubation at 30°C with shaking, diluted aliquots of the culture were plated on LB medium to estimate the titer of the population and transformants were scored on selective media. Transformation frequencies were calculated as transformants per recipient.

### Evolution experiments

The plasmid allele dynamics evolution experiment was conducted with ancestral plasmid carrying *A. baylyi* populations under two selection regimes, three novel initial allele frequencies and two population bottleneck sizes. The ancestral plasmid present was pTAD-R for all populations (n=72). On the onset of the experiment, *A. baylyi* strain BD413 containing the ancestral plasmid was naturally transformed with the donor DNA following the previously described protocol [28]. Donor DNA was added at specific concentrations to generate diverse transformants frequencies. Three different initial allele frequencies were created: 10^−4^ (700ng/ml donor DNA), 10^−6^ (7ng/ml donor DNA) and 10^−7^ (0.7ng/ml donor DNA). After the transformation was complete (i.e., after 90 minutes of incubation), a fraction of the cells was plated to determine the total cell population and the transformation frequency. The viable titer of the cultures was obtained from the cells plated on LB media (10^−4^, 10^−5^ and 10^−6^ dilutions), whereas the transformation efficiency was calculated by plating on LB + kanamycin (5 μg/ml) (1, 10^−1^ and 10^−2^ dilutions). Cells that had restored the *nptII* gene on one of copies of the recipient plasmid survived the kanamycin treatment. After overnight growth cultures were diluted into fresh medium with 1:100 (weak bottleneck) or 1:1000 (strong bottleneck) dilution factor. The bacterial populations were subjected to two selective regimes, non-selective conditions and selective conditions (kanamycin 0.375 μg/ml). The cultures were incubated in 96-deep-well plates with a total volume of 1 ml at 30 °C with constant shaking. The transfer regime was then maintained for 30 transfers, which corresponds to approx. 200-300 generations (~6-7 generations per transfer in liquid medium for the weak bottleneck and ~9-10 generations for the strong bottleneck). Every day the population sizes were estimated, the total population size was estimated from the plating on LB media and the proportion of hosts in the population was measured by plating on selective media, LB supplemented with kanamycin. The distinction between heterozygotes and homozygotes for the new allele (*nptII*) comes from the gene of the reporter construct. For our model plasmid, the fluorescence that confers the GFP protein enables us to distinguish them.

The chromosomal allele experiment was performed by employing a mixture of two *A. baylyi* strains, BD4 (tryptophan prototroph) carrying a *tetA* gene which is the ancestral allele in this setting and BD413 (tryptophan auxotroph) carrying a *nptII* gene which is the novel allele. The strains were then mixed in three ratios comparable to the initial novel allele frequencies of the plasmid experiment, BD4/BD413 at 10^−4^, 10^−6^ and 10^−7^ proportions. A fraction of the cell was plated to determine the initial host proportion and the cell titers were calculated as previously described employing M9 minimal media and LB supplemented with kanamycin (5 μg/ml) as selective media to distinguish both strains. The experimental set up was identical as the one employed for the plasmid allele dynamics experiment, with serially passaged populations employing a 1:100 or 1:1000 dilution factor under non-selective and selective conditions (n=72). The bacterial populations were subjected to two selective regimes, non-selective conditions and selective conditions (kanamycin 0.4 μg/ml). The cultures were transferred for 30 transfers and population size estimation was done for each day.

### Statistical analysis

All statistical tests and data analysis were performed in RStudio version 1.2.1327. Comparisons between groups (e.g., fitness calculations) were performed with two sided Wilcoxon signed-rank test [40] by the “wilcox.test” function.

### Simulation

We simulate the population dynamics of the serial dilution experiment using a random segregation model for multicopy replicons(27,29], which we extended for periodic bottlenecks. The model comprises competitive growth of bacteria, periodic bottlenecks, and plasmid segregation. In our model, cells with at least one copy (*i* ≥ 1) of the novel (resistance-conferring) plasmid variant reproduce at an intrinsic per-capita rate *r*_i_ = 1, where *j* denotes the cell type. Bacteria without the novel plasmid variant reproduce at a rate *r*_0_ = 1 − *s*, where *s* > 0 reflecting an antibiotic environment or *s* = 0 for a neutral environment. At the beginning of the simulation, the population consists of *<u>N</u>*^(0)^ = 10^7^ cells with a (resistant) subpopulation of proportion *f*_0_ that carries one (out of *n*_rc_) replicon copy of the novel variant. All cells have a fixed copy number of the plasmid *n*_rc_ at cell birth. We describe two modes of plasmid replication during cell division, regular and random replication. For regular replication, all plasmid copies are duplicated in a parental cell before it divides. For random replication, plasmid replication follows a successive manner. First, one plasmid is randomly chosen from the plasmid pool for replication and the two replicates are added to the pool of plasmids (see also [33]). This procedure is repeated *n*_rc_ - 1 more times such that a cell – as for regular replication – carries 2*n*_rc_ plasmid copies before cell division. Finally, plasmid copies are distributed to the bacterial daughter cells at random, but in equal numbers to comply with the fixed plasmid copy number *n*_rc_ [cf. 27,29]. Bacterial populations grow up to a carrying capacity 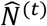 before the transfer *t* occurs, which we determine separately for each individual transfer and condition as the mean over the six experimental replicates. We study the deterministic dynamics of bacterial growth arising from large population sizes, which can be computed by integrating a system of *n* + 1 ordinary differential equations until the corresponding carrying capacity is reached. The differential equations describes the temporal change of the cell numbers *N*_*i*_ for the individual cell types *i*=0,…,*n*_rc_ (cells carrying *i* novel plasmid copies) between two bottlenecks, *t-1* and *t*, caused by growth and random segregation of plasmid variants at cell divisions. The time derivative for any cell type *i* is given by

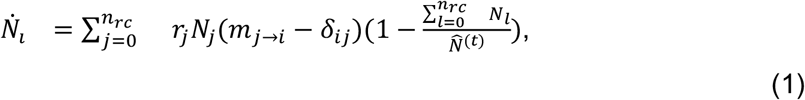

where *m*_*j*→*i*_ denotes the expected number of *i*-type cells produced at division of an *j*-type cell and *δ*_*ij*_ denotes Kronecker’s delta (i.e., *δ*_*ij*_ = 1 for *i* = *j* and *δ*_*ij*_ = 0 otherwise). For regular replication, we have

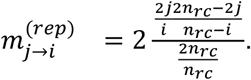

In case of random replication, we have

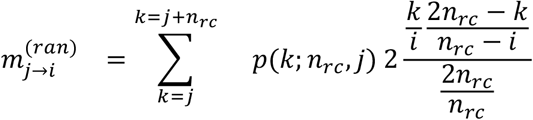

[see 29]. *p*(*k*; *n*_rc_, *j*) is the probability to have *k* mutated copies before cell division if plasmid copies are replicated in a successive manner (see random replication above). For heterozygous cells, 0 < *j* < *n*_rc_, the probability *p*(*k*; *n*_rc_, *j*) is given by the shifted probability mass function of a beta-binomial distribution, 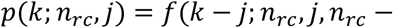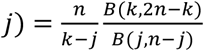 with the Beta-function 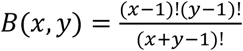. For homozygous cells, *j* = 0 or *j* = *n*, we have *p*(*k*; *n*_rc_, *j*) = *δ*_3,9)_. We define the end of the growth phase by the time when the time derivative of the various cell numbers, *N_l_*, becomes very low, |*Ṅ_l_*| ≤ 0.01 for all cell types *i*, i.e., the change in the cell number of any cell type is at maximum 1% of the per-capita growth rate of wild-type cells At this time, the carrying capacity is almost reached as the change in the total cell number 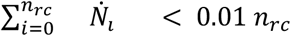 becomes very low. At the end of the growth phase, populations undergo a bottleneck of factor *b*, which we model stochastically by drawing a proportion *b* of the population at carrying capacity (rounded to an integer number) using a multinomial distribution generated by the proportions of all cell types *i* in the population.

### Estimation of model parameter of selection *s*

We used optimization methods to estimate the model parameter ***s*** from the time series of kanamycin resistant cell frequencies. Optimal *s* values were obtained by performing a nonlinear least-square analysis with the residuals

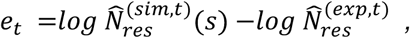

where 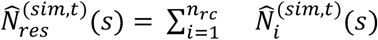 and 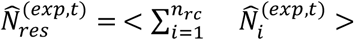 denotes the resistant cell frequencies before transfer *t* in computer simulations for a given parameter *s* and in the experiment respectively [41]. We transformed simulation and experimental data using a logarithmic scale due to the exponential growth in our model (see Eq. [1]). We use the conditions with high initial frequencies *f*_0_ = 10-4 and weak bottlenecks *b* = 10-2 for the parameter estimation to minimize the stochasticity from population bottlenecks. The experimental data was averaged over the six experimental replicates indicated by the angle brackets in the latter equation. For the computer simulation, we use only single trajectories for the optimization due to the low variability under these conditions (see Fig 3a, plot 1 and 3). The nonlinear regression to estimate *s* was performed using the Nelder-Mead method implemented in the Python package *lmfit (*function *minimize*, parameter *method=’nelder’*).

## Data Availability

The source data underlying Figs 1 and 2 and Figs S1, S2, S3, S4, S5 and S6 are provided as a Source Data file. The code for our simulations is available online (https://github.com/mariosanter/multicopy-bottle-fixation).

## Acknowledgments

We thank Daniela Kluger and Ian Dewan for their assistance in the experimental work. The authors thank Yiqing Wang, Shreya Vichare and Ishan Bhatt for critical comments on the manuscript, the entire Genomic Microbiology group and Stochastic Evolutionary Dynamics group for their help and discussions. AG is supported by the International Max Planck Research School (IMPRS) for Evolutionary Biology and by the Leibniz Science Campus EvoLUNG. MS is a member of the International Max Planck Research School for Evolutionary Biology and gratefully acknowledges the benefits provided by the program. This work was funded by the Deutsche Forschungsgemeinschaft (DFG, German Research Foundation) - project number 418432175 (HU).

## Author Contributions

AG, MS, NFH, HU and TD conceived the study and designed the experiments. AG established and performed the experimental work with the help of NFH. MS performed the mathematical modelling and the simulations. NFH, HU and TD supported the data analysis. All authors interpreted the results and wrote the manuscript.

## Supporting information

**S1 Fig.**
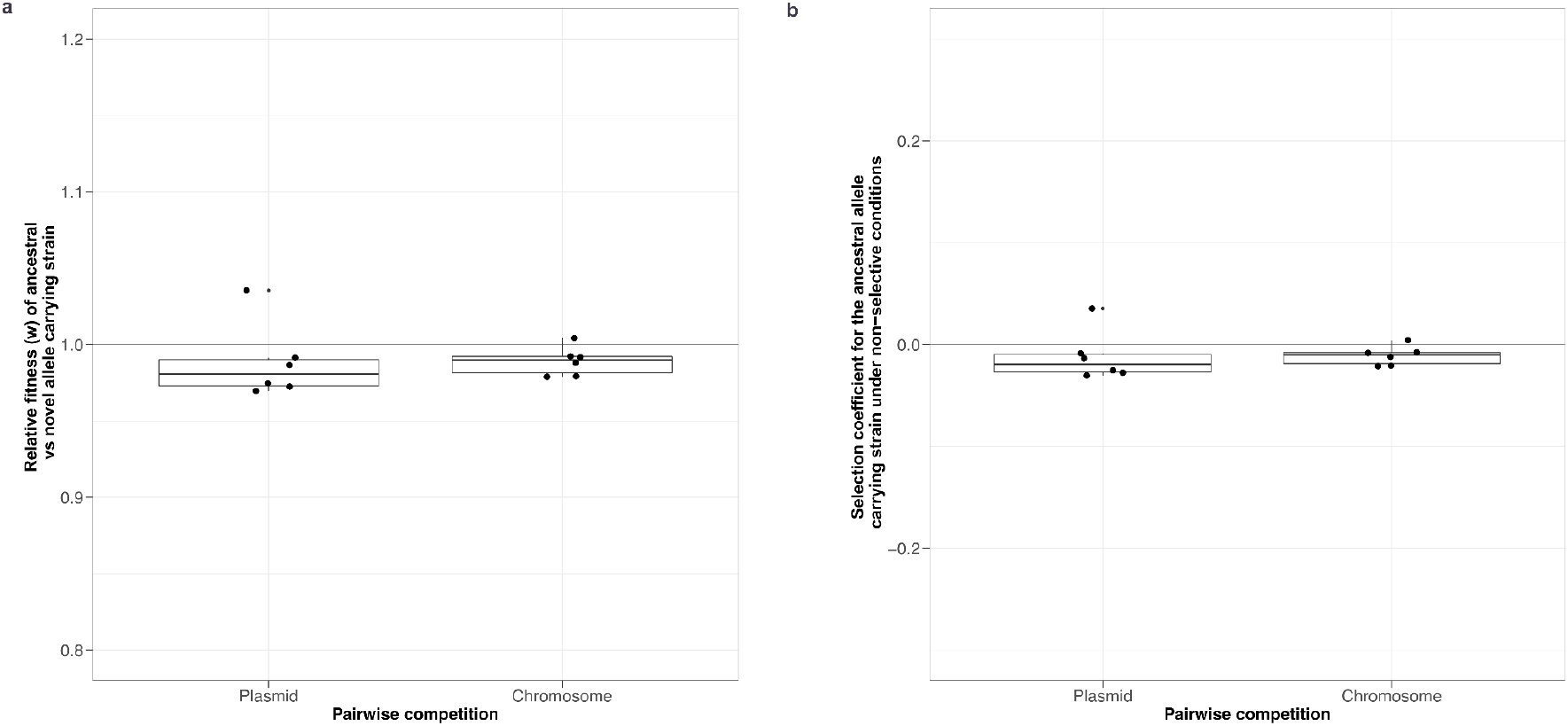
Competition experiments to quantify the fitness differences between the ancestral and novel alleles homozygotes. a) Pairwise competition experiments between plasmid carrying strains *A. baylyi* BD413 pTAD-R (*GFP*, ancestral plasmid allele) and *A. baylyi* BD413 pTAD-R (*nptII*, novel plasmid allele); *A. baylyi* BD4 (*tetA*, ancestral chromosome allele) and *A. baylyi* BD413 (*nptII*, novel chromosomal allele). The experiments were conducted under non-selective conditions for 120h at 30°C with 6 biological replicates per replicon. Fitness was calculated relative to the growth as described in refs. [43,44], no significant fitness effect could be observed (pTAD-R H_0_: w = 1, p = 0.4375, n = 6 & Chromosome H_0_: w = 1, p = 0.0625, n = 6. Wilcoxon test). b) Selection coefficient applicable to the ancestral allele carrying strain under non-selective conditions. Selection coefficient was calculated by the formula s = w-1 [43]. No significant selection coefficient was applicable to any strain.

**S2 Fig.**
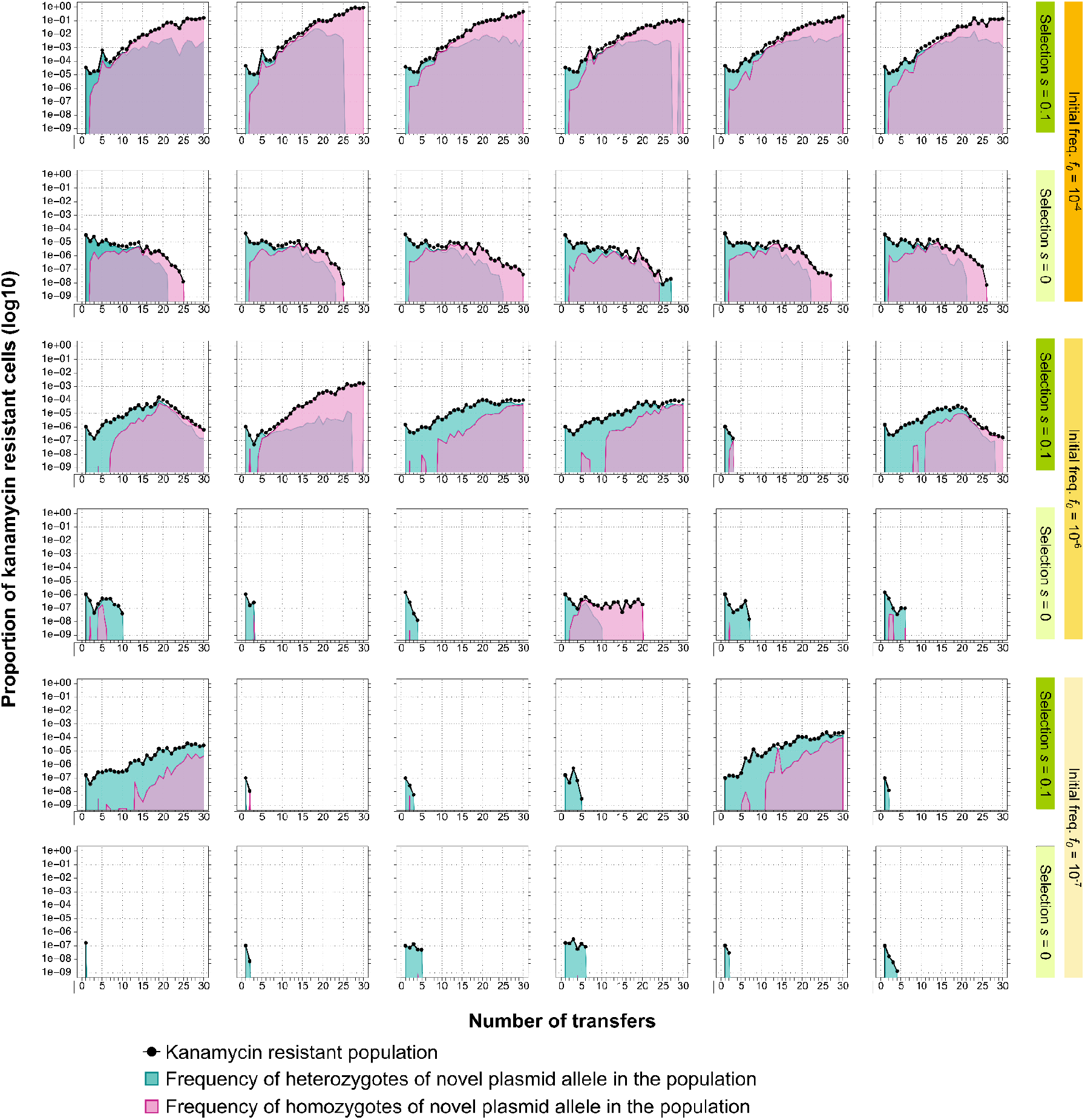
Plasmid allele dynamics in nonselective and selective conditions under weak bottleneck (1:100 or 10^−2^). (A) The dynamics of pTAD-R over approximately 200 generations. The novel allele was introduced at the beginning of the experiment employing different donor DNA concentration to achieve 3 initial allele frequencies. Host frequencies in the total population are shown in black, heterozygote frequencies are shown in green and novel homozygotes in pink. a) High initial frequencies b) Moderate initial frequencies c) Small initial frequencies

**Fig S3.**
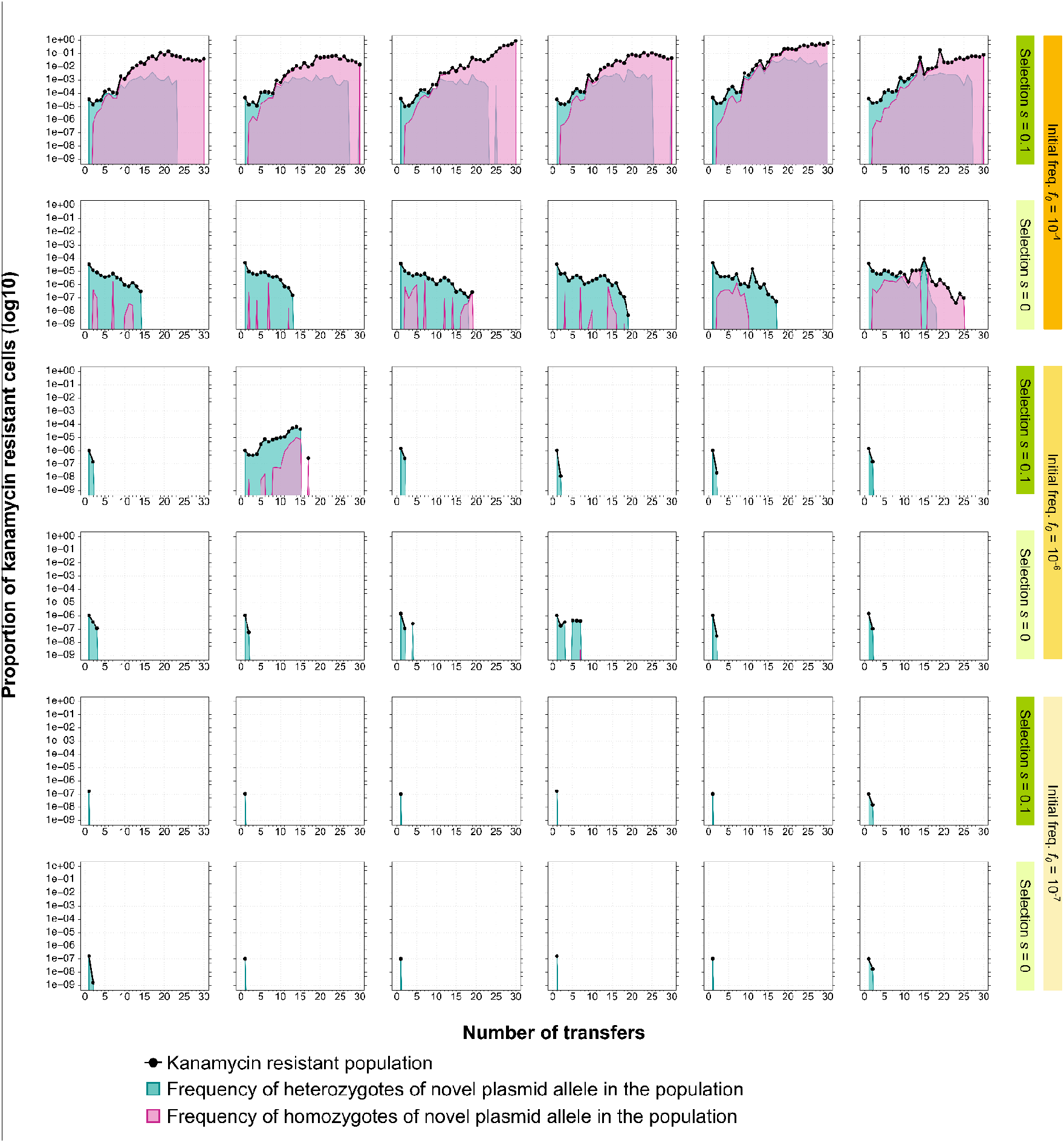
Plasmid allele dynamics in nonselective and selective conditions under strong bottleneck (1:1000 or 10^−3^). (A) The dynamics of pTAD-R over approximately 200 generations. The novel allele was introduced at the beginning of the experiment employing different donor DNA concentration to achieve 3 initial allele frequencies. Host frequencies in the total population are shown in black, heterozygote frequencies are shown in green and novel homozygotes in pink. a) High initial frequencies b) Moderate initial frequencies c) Small initial frequencies

**S4 Fig.**
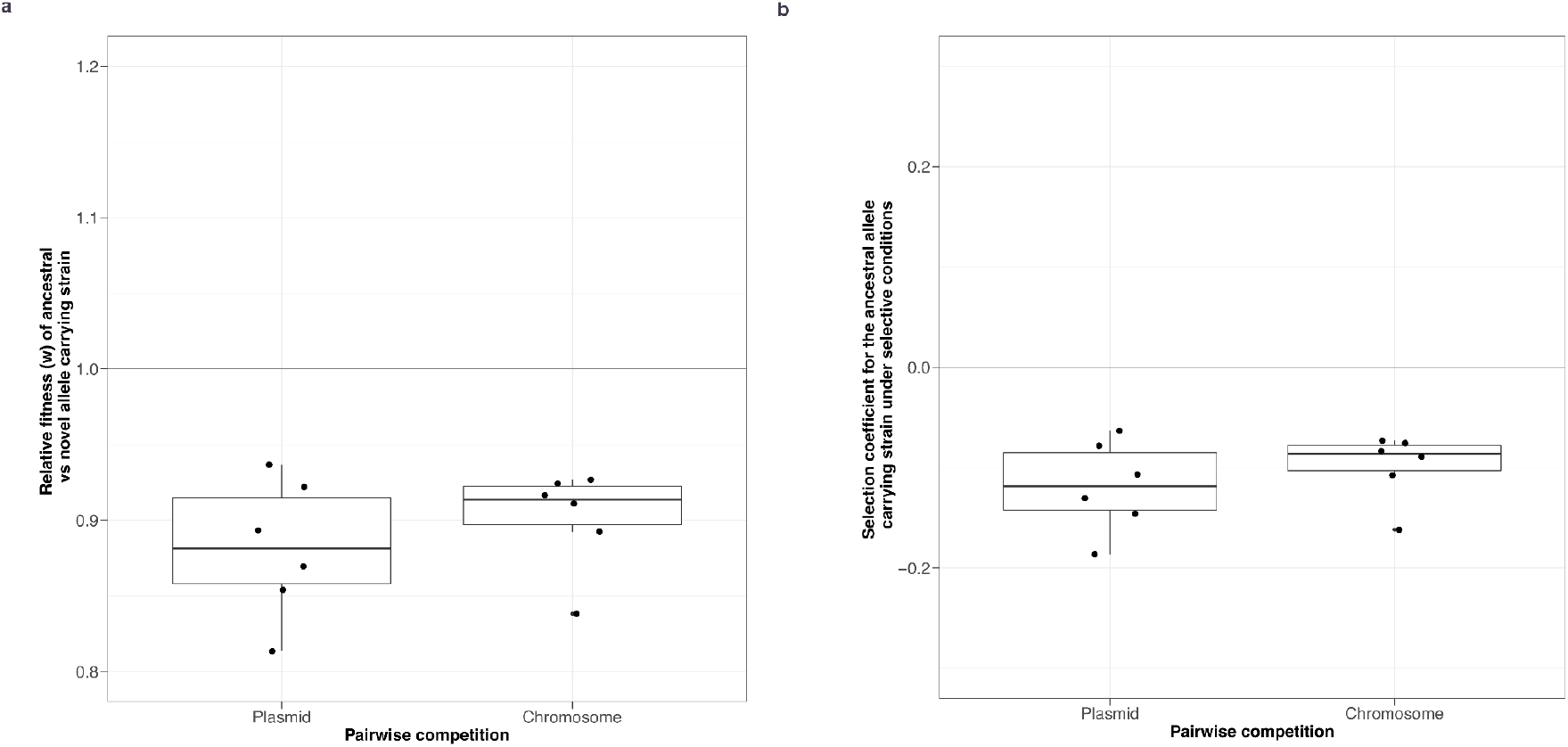
Competition experiments to quantify the fitness differences between the ancestral and novel alleles homozygotes under selective conditions. a) Pairwise competition experiments between plasmid carrying strains *A. baylyi* BD413 pTAD-R (*GFP*, ancestral plasmid allele) and *A. baylyi* BD413 pTAD-R (*nptII*, novel plasmid allele); *A. baylyi* BD4 (*tetA*, ancestral chromosomal allele) and *A. baylyi* BD413 (*nptII*, novel chromosomal allele). The experiments were conducted under selective conditions (kanamycin_plasmid_=0.375μg/ml and kanamycin_chromosome_=0.4μg/ml) for 120h at 30°C with 6 biological replicates per replicon. Fitness was calculated relative to the growth as described in refs. [43,44], a significant fitness effect could be observed (pTAD-R H_0_: w = 0.9, p = 0.4375, n = 6 & Chromosome H_0_: w = 0.9, p = 5625, n = 6. Wilcoxon test). b) Selection coefficient applicable to the ancestral allele carrying strain under selective conditions. Selection coefficient was calculated by the formula s = w-1 [43]. A significant selection coefficient was applicable to the ancestral strain. Ancestral allele containing strain shows a selective disadvantage (s<0) of 0.1.

**S5 Fig.**
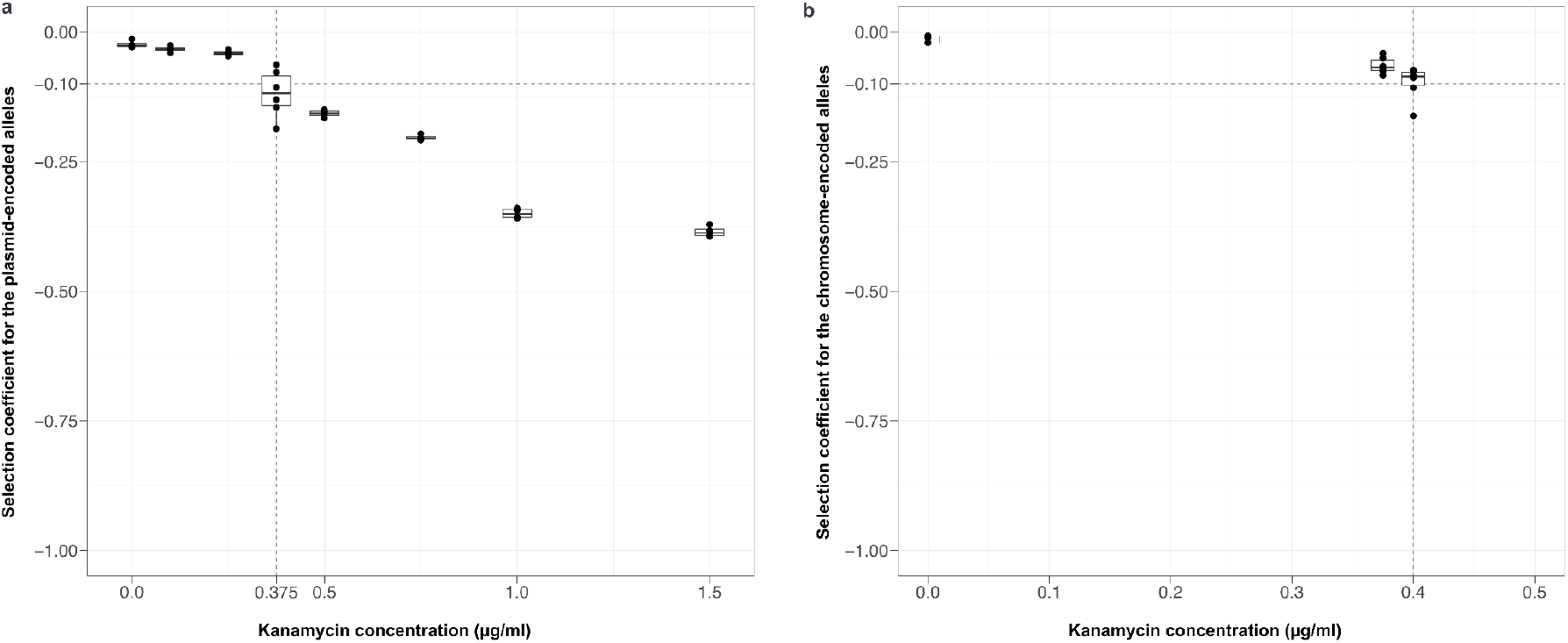
Competition experiments to quantify the fitness differences between the ancestral and novel alleles homozygotes under diverse selective conditions. a) Pairwise competition experiments between plasmid carrying strains *A. baylyi* BD413 pTAD-R (*GFP*, ancestral plasmid allele) and *A. baylyi* BD413 pTAD-R (*nptII*, novel plasmid allele); *A. baylyi* BD4 (*tetA*, ancestral chromosomal allele) and *A. baylyi* BD413 (*nptII*, novel chromosomal allele). The experiments were conducted under diverse selective conditions (kanamycin_plasmid_=0-1.5μg/ml and kanamycin_chromosome_=0-0.4μg/ml) for 120h at 30°C with 4-6 biological replicates per kanamycin concentration assayed. Fitness was calculated relative to the growth as described in refs. [43,44]. Selection coefficient applicable to the ancestral allele carrying strain under selective conditions. Selection coefficient was calculated by the formula s = w-1 [43].

**S6 Fig.**
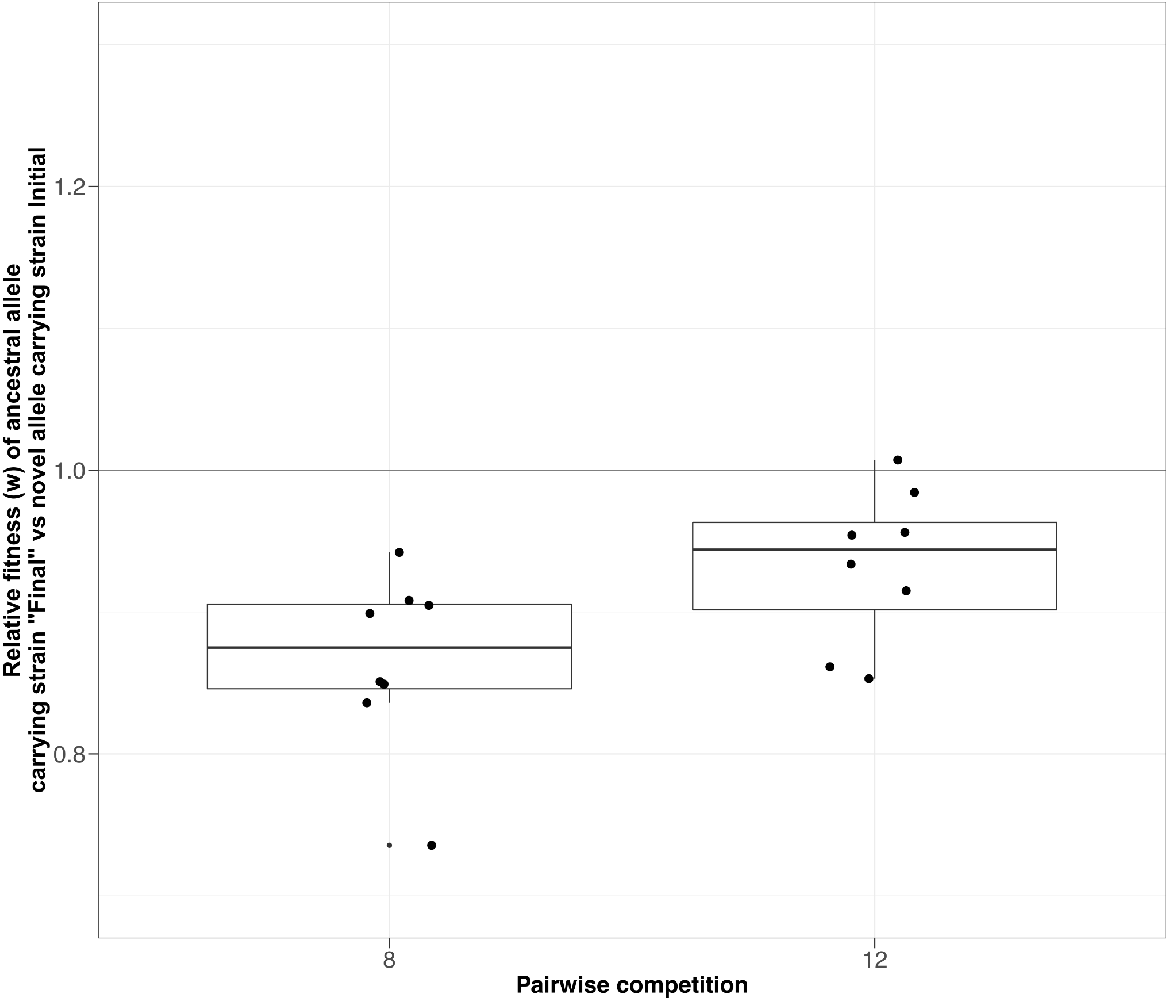
Quantification of fitness differences under selective conditions between non-host population at the end of the experiment and the host population at the beginning of the experiment. Pairwise competition experiments between host and non-host plasmid populations of experimental conditions 8 and 12. The experiments were conducted in selective medium (kanamycin 0.375 μg/ml) for 120h at 30°C with 8 biological replicates per condition. Fitness was calculated relative to the growth as described in refs. [43,44]. A significant fitness disadvantage of the non-host population could be observed (Cond 8 H_0_: w>1, p= 0.003906, n=8 & Cond 12 H_0_: w>1, p= 0.007813, n=8). Data also shows a fitness advantage of the host over the non-host population with a selection coefficient of 0.1 for the non-hosts (Cond 8 H_0_: w=0.9, p= 0.25, n=8 & Cond 12 H_0_: w=0.9, p= 0.1484, n=8).

**S7 Fig.**
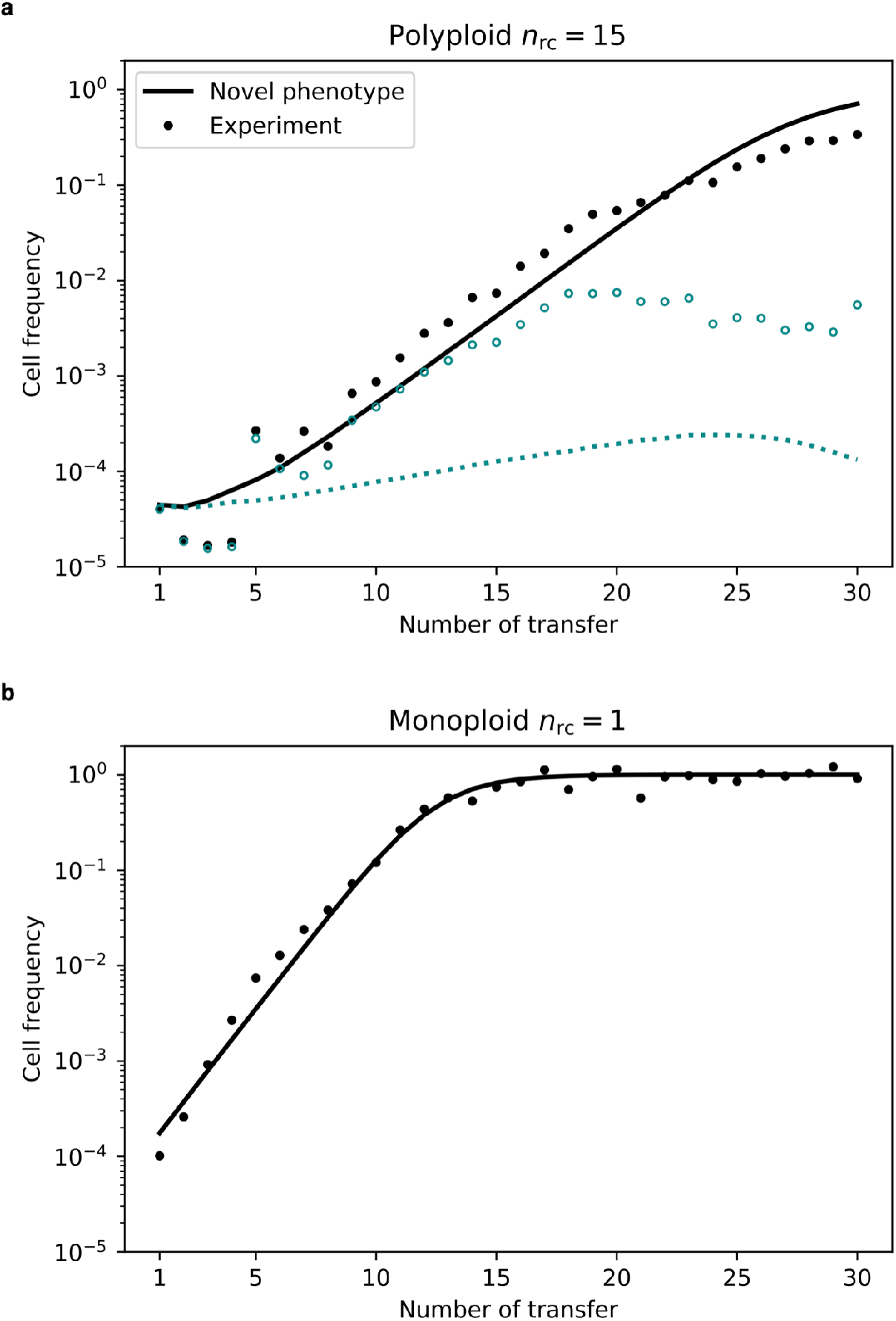
Comparison of simulated allele dynamics with *s* values yielding an optimal fit with experimental results. a) The simulated allele dynamics on the polyploid replicon (*n*_rc_ = 15) with a selection coefficient *s* = 0.086, which yields an optimal fit to the experimental allele dynamics on the chromosome. b) The simulated allele dynamics on the monoploid replicon (*n*_rc_ = 1) with a selection coefficient *s* = 0.14, which yields an optimal fit to the experimental allele dynamics on the multicopy plasmid. We fitted the selection coefficients using the cell frequencies of the novel phenotype (kanamycin resistant) for the experimental conditions *f*_0_ = 10-4 and *b* = 0.01.

**S8 Fig.**
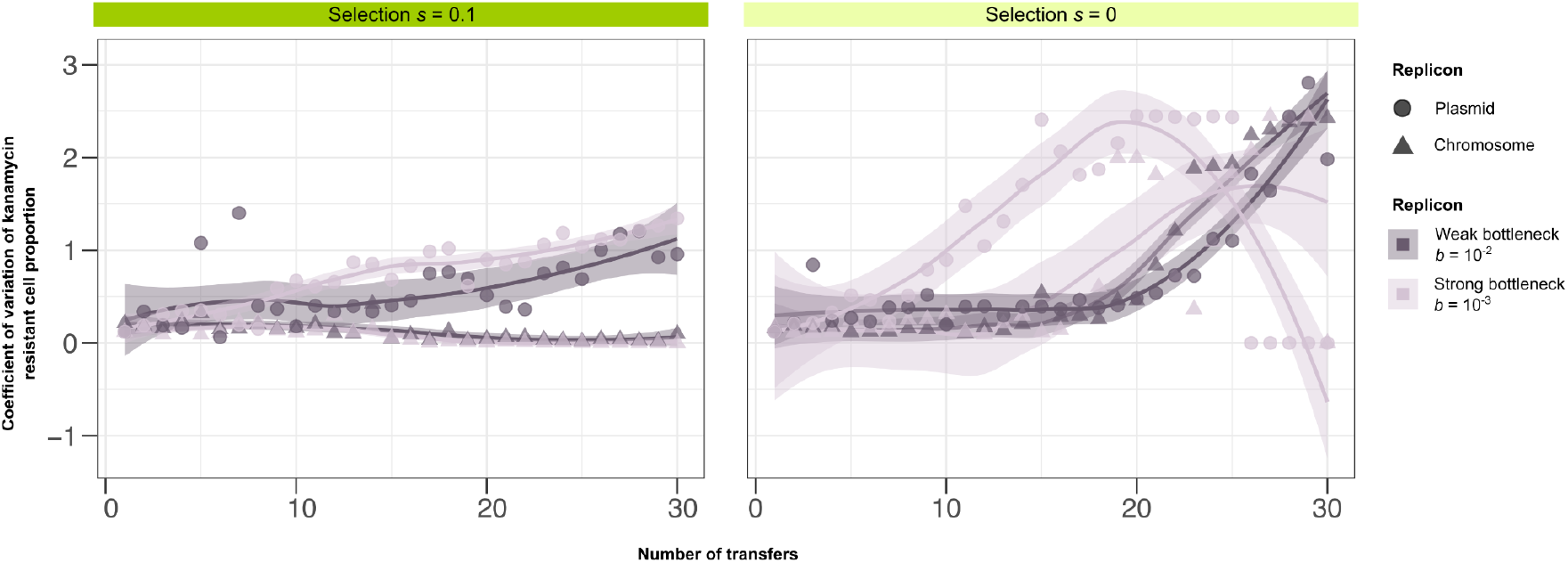
Coefficient of variation of kanamycin resistant cell proportion for high initial allele frequency populations (*f_0_* = 10^−4^). Lines present the trendline and the shaded area corresponds to the confidence interval.

**S1 Table.**
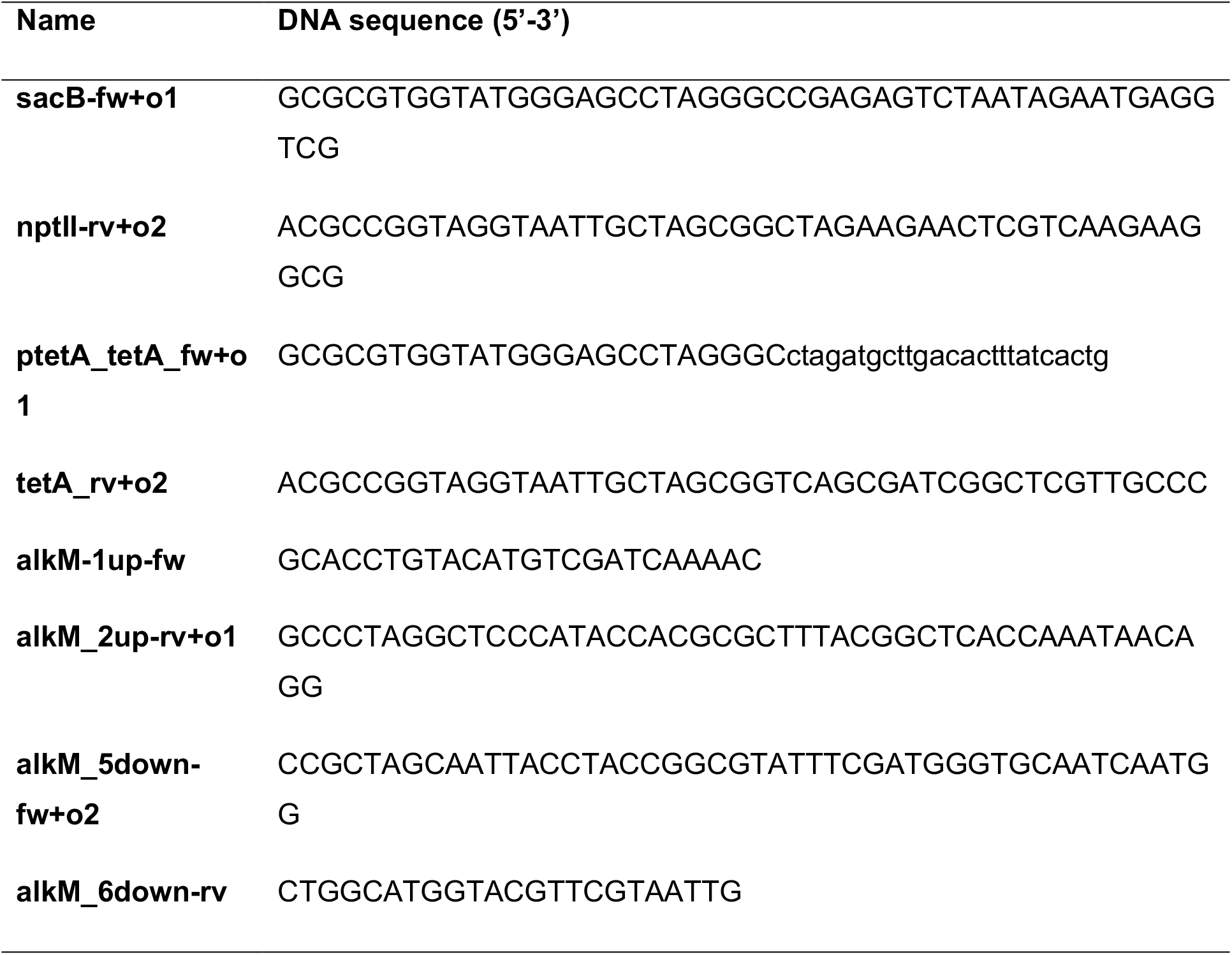
Primers used in the study.

